# The Physical Biology of Nucleolus Disassembly

**DOI:** 10.1101/2023.09.27.559731

**Authors:** An T. Pham, Madhav Mani, Xiaozhong A. Wang, Reza Vafabakhsh

## Abstract

During cell division, precise and regulated distribution of cellular material between daughter cells is a critical step and is governed by complex biochemical and biophysical mechanisms. To achieve this, membraneless organelles and condensates often require complete disassembly during mitosis. The biophysical principles governing the disassembly of condensates remain poorly understood. Here, we used a physical biology approach to study how physical and material properties of the nucleolus, a prominent nuclear membraneless organelle in eukaryotic cells, change during mitosis and across different scales. We found that nucleolus disassembly proceeds continuously through two distinct phases with a slow and reversible preparatory phase followed by a rapid irreversible phase that was concurrent with the nuclear envelope breakdown. We measured microscopic properties of nucleolar material including effective diffusion rates and binding affinities as well as key macroscopic properties of surface tension and bending rigidity. By incorporating these measurements into the framework of critical phenomena, we found evidence that near mitosis surface tension displays a power-law behavior as a function of biochemically modulated interaction strength. This two-step disassembly mechanism, which maintains structural and functional stability of nucleolus while allowing for its rapid and efficient disassembly in response to cell cycle cues, may be a universal design principle for the disassembly of other biomolecular condensates.

## Main Text

Precise and regulated distribution of cellular material between daughter cells during cell division is a critical step (1-3). Consequently, many cellular strategies and cell cycle-dependent checkpoints have evolved to ensure fidelity in partitioning of genomic and non-genomic cellular components (4-6). For membrane-bound organelles such as the nuclear envelope, endoplasmic reticulum, mitochondria, endosomes, and the Golgi apparatus division relies on remodeling or fragmentation of organelles into many membrane-bound units (2, 3, 7-9). These units disperse throughout the cytoplasm, either passively or through interactions with the cytoskeleton, to segregate and eventually divide between daughter cells, and sometimes with a degree of stochasticity (10, 11). In contrast, division of condensates and membraneless organelles requires a distinct approach due to their potentially low abundance and the fact that they are unable to undergo fragmentation into membrane-delimited units that encapsulate their contents and subsequently re-fuse after mitosis (12-15). The specific regulatory mechanisms and checkpoints responsible for the accurate division of condensate material while ensuring the synchronization of condensate disassembly with the cell cycle cues remains poorly understood.

The interphase nucleolus in eukaryotic cells is the largest membraneless nuclear organelle and serves as the primary site for the ribosomal RNA (rRNA) transcription and processing, and ribosome assembly in eukaryotic cells (16, 17). Additionally, it is a key contributor to stress sensing, cellular proliferation, and maintenance of genome organization (17, 18). During mitosis, nucleoli undergo complete disassembly followed by rapid reassembly at the end of mitosis which enables the dispersion and distribution of nucleolar material between daughter cells with high fidelity (12, 13, 16). The disassembly process depends on phosphorylation of RNA polymerase I (Pol I) specific transcription factors and other nucleolar proteins by cell cycle-dependent kinases such as cyclin-dependent kinase 1 (CDK1) which affect their enzymatic activity or binding affinities (19, 20). Therefore, continuous processing of rRNA and precise and finely tuned affinity between nucleolar components is important for nucleolus function (16, 21). Interruption of these factors results affect nucleolus structure and function (22, 23). These findings suggest that there is a direct correlation between the biochemical activity of the nucleolus and its biophysical and material properties. Despite advances in our understanding of the biochemical composition of the nucleolus (24, 25), and the discovery of liquid-liquid phase separation (LLPS) as a general mechanism underlying its organization (15, 18, 26), it remains unclear how the macroscopic biophysical features of the nucleolus are related to the properties of nucleolus material at the microscopic scale, and the mechanisms that underlies their correlated dynamics during the disassembly process. In complex systems the connection between microscopic and macroscopic features as a system approaches a critical regime has been studied extensively (27-29). Molecular dynamics simulation of condensates made of an ideal multivalent polymer that undergoes temperature change has shown that the framework of critical phenomena can be used to estimate thermodynamic parameters from experimental observables in such systems (30). On the other hand, cellular functions operate at a constant temperature, which presents a challenge when attempting to contextualize the process of nucleolus disassembly during mitosis within the framework of critical phenomena. Here we report a physically inspired strategy that leverages the information derived from nucleolar shape fluctuations, molecular binding kinetics, and bulk properties of the nucleolus to uncover the underlying principles governing its disassembly and its coordination with cell division.

## Results

We generated a HeLa cell line stably expressing mClover3-H2B (to visualize the DNA) and nucleophosmin 1(NPM1)-mRuby3 (to visualize the nucleolus) and used 4D confocal live-cell imaging to continuously measure the bulk structural dynamics of nucleolus through the cell cycle (Fig. 1A, Fig. S1). NPM1 is a highly abundant protein in the granular component (GC) of nucleolus with a small fraction also present in the nucleoplasm (19). Since NPM1 is a key driver for formation of nucleolus it has been used as an informative marker for tracking the nucleolus dynamics (31-33). We visually picked the cells at early prophase based on the appearance of chromatin and imaged them as they progressed through mitosis. We monitored the dynamics of nucleolar disassembly by tracking and quantifying the distribution of NPM1 proteins during mitosis.

**Figure 1.**
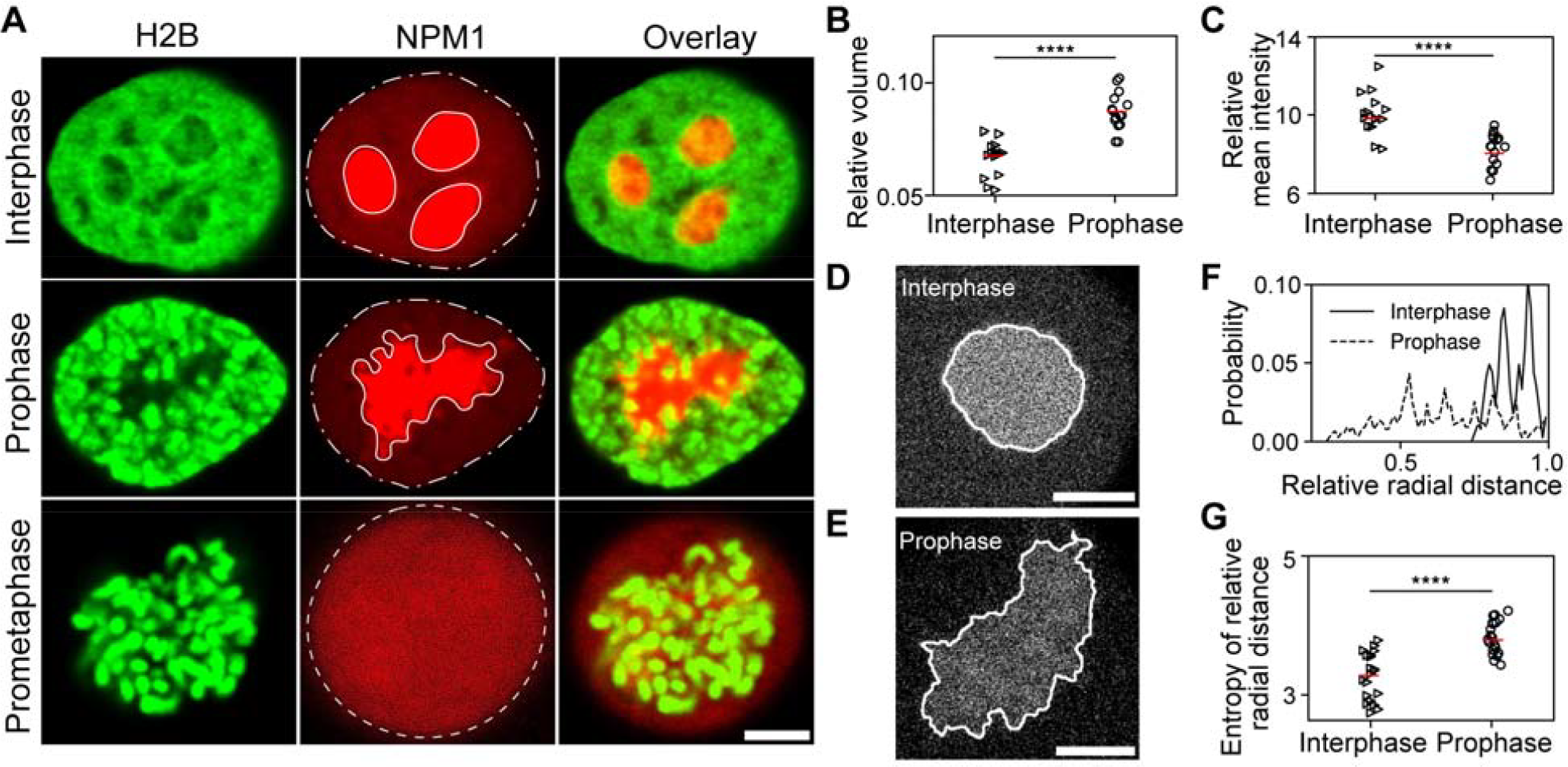
Morphology changes of nucleolus at mitosis entry. (**A**) Live HeLa cells expressing mClover3-H2B and mRuby3-NPM1 in interphase and mitosis. White dash-dot and solid lines outline segmented nucleus and nucleolus regions for feature quantification. White dashed line shows cell membrane boundary after nuclear membrane and nucleolus disassemble at prometaphase. (**B**) Relative volume (volume of nucleolus/volume of nucleus) and (**C**) relative mean intensity (mean intensity of nucleolus/mean intensity of nucleus) in interphase (triangle) and prophase (circle) cells. Live HeLa cells expressing mRuby3-NPM1 in interphase (**D**) and prophase (**E**). White solid lines represent interfacial boundaries between nucleolus and nucleoplasm. (**F**) Distribution of the relative radial distance in interphase and prophase cells. (**G**) Entropy of relative radial distance measurement for interphase (triangle) and prophase (circle). For all boxplots (B, C, G), red bars indicate the mean. Significance was tested by t-test (****P<0.0001). Biological replicates: n = 15 cells (B, C). n = 20 cells (G); Scale bar, 5 μm (A); 3 μm (D, E).

### Nucleolar morphology begins to change upon cell entry into mitosis

We first developed an image analysis pipeline to quantify the change in the nucleolar volume and the nucleolar NPM1 concentration as cells entered prophase (Supplemental Discussion, Fig. S2). We found that the normalized nucleolus volume was larger for cells in the prophase compared to cells in the interphase (Fig. 1B). Consistently, we observed that the normalized nucleolar NPM1 density was lower in the prophase cells (Fig. 1C). These data suggest that nucleoli undergo global structural changes as cells progress in cell cycle towards mitosis.

To quantify nucleolar shape change between prophase and interphase cells (Fig. 1A), we extracted nucleolar perimeter at its equatorial plane (Figs. 1D, 1E and Supplemental Discussion) and plotted the normalized radial distance of all nucleolar boundary positions (Fig. 1F) and the entropy of radial distances (Fig. 1G) as shape-descriptive features (34). Consistent with our qualitative observation, this analysis showed that the nucleoli of cells in interphase were more round with smooth boundaries while the nucleoli of cells in prophase were morphologically more irregular in shape. The entropy of the radial distance distribution also exhibited a significantly larger geometric entropy for cells in prophase relative to the interphase (Fig. 1G). We hypothesized that the observed geometric difference in the shape of the nucleolus might be indicative of a structural instability as cells enter prophase, which in turn could drive the loss of integrity of the nucleolus.

### Nucleolus dissembles in two steps

To gain insights into the dynamic nature of nucleolus disassembly throughout the mitosis and to determine whether the observed change is a continuous process or segmented into discrete steps, we collected volumetric images of HeLa cells from prophase to mitosis (Fig. 2A, Movie S1). We observed an abrupt transition from a state of two-phase coexistence between a dense phase and a dilute phase at the prophase to a single homogeneous phase immediately following nuclear envelope breakdown (Fig. 2A, time point zero). To quantify this process, we developed a novel analysis platform based on measuring the empirical probability density function (PDF) of pixel intensities at each frame. Notably, this analysis eliminates the need for a threshold parameter, crucial given the drastic temporal changes in pixel intensity distribution. With this method, we quantified the disassembly trajectory as cells progressed towards mitosis (Fig. 2B, Fig. S3). We fitted the empirical distributions with double power-law functions, corresponding to a dense and a dilute phase, respectively (Fig. S3A).

**Figure 2.**
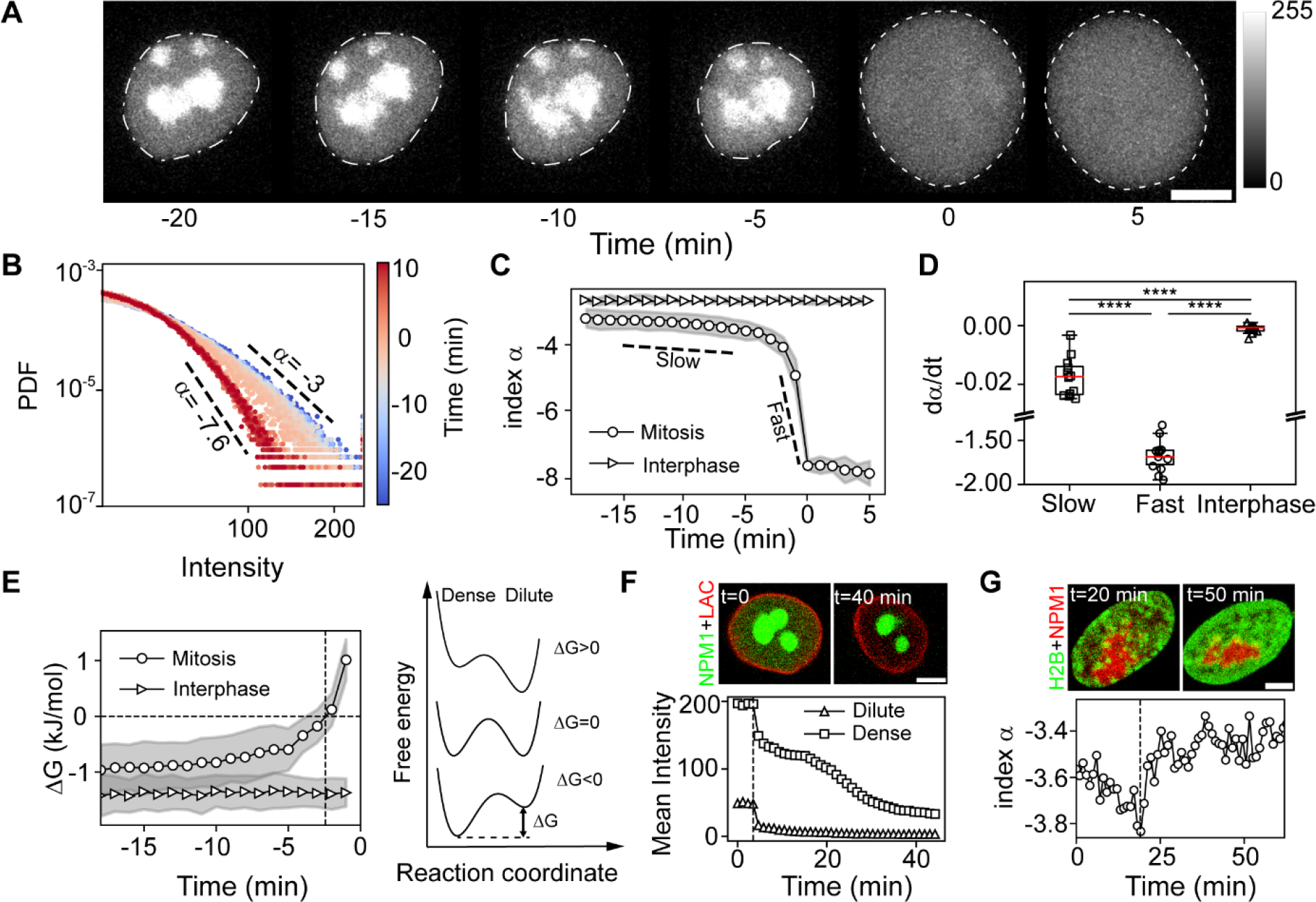
Dynamics dissociation of nucleolus. (**A**) Time-lapse of HeLa cell expressing mRuby3-NPM1 progressing from prophase to prometaphase. Dash-dot lines denote nucleus, dashed lines illustrate cell membrane boundary. t = 0 denotes nuclear envelope breakdown. (**B**) Probability density function of pixel intensities at different time points at 1 minute time intervals. t = 0 denotes the time of nuclear envelope breakdown. Dashed lines represent the power law exponent coefficients (α) of dense phase in log-log scale. (**C**) Power law exponent coefficients (α) as a function of time for cells in interphase and mitosis. n=12 cells. Slow and fast slopes (dashed lines) highlight two phases of the nucleolus disassembly. (**D**) Slow and fast disassembly rates for cells mitosis and interphase. Red bars indicate the mean and box extend from the first to third quartile. Significance was tested by t-test (****P<0.0001). (**E**) Left, NPM1 free energy of transfer (ΔG) from nucleoplasm to nucleolus as a function of time for cells in mitosis and interphase. n = 12 cells. Dash vertical line indicates time when the free energy becomes positive. Right, schematic showing changes of free energy landscape as NPM1 relocated from dense to dilute phases through mitosis. (**F**) Top, representative images of HeLa cells expressing EGFP-NPM1 and mRuby2-Lamin A/C before (t=0 min) and after (t=40 min) Triton-X 100 treatment at t = 5 min. Bottom, quantification of mean intensity of NPM1 in dense and dilute phases. (**G**) Top, representative images of HeLa cells expressing mClover3-H2B and mRuby3-NPM1 right before (t=20 min) and after (t=50 min) RO-3306 treatment. Bottom, quantification of the nucleolus disassembly before and after treatment. Dash vertical line indicates time adding Triton-X 100 (F) and RO-3306 (G). Data points and shaded areas (C, E) represent mean ± s.d. Scale bar, 8 μm (A); 5 μm (F, G).

Quantitatively, the slope of the dense phase (α) varied continuously as cells progressed towards nuclear envelope breakdown, rendering it an effective index to track the dynamics of the disassembly process (Fig. 2B). Consistent with the transfer of material from a dense to a dilute phase, α decreased slowly during early prophase, followed by a steep drop at the nuclear envelope breakdown (Fig. 2C). During the rapid second phase, the rate of change of α increased by two orders of magnitude compared to the slow phase (Fig. 2D). Importantly, interphase cells showed only minor decreases over a similar imaging window (Fig. S3B), indicating that the nucleolus is stable during interphase and the measured slow phase is not an artifact of photobleaching (Figs. 2C, 2D).

To align the timing of nucleolus disappearance at the end of the fast phase with the cell cycle stage of cells, we performed an experiment with a new HeLa cell line that stably expressed mRuby2-Lamin A/C and EGFP-NPM1 (Fig. S4A). In this cell line we observed that the nucleolus disappearance coincided with the complete disappearance of the nuclear envelope (Fig. S4B, Movie S2). Therefore, the fast phase of nucleolus disassembly coincides with nuclear envelope breakdown and subsequent dilution of the nuclear material.

The above analysis of pixel intensities allowed us to systematically identify the intensity threshold separating the dilute and the dense phases (Fig. S3A), as well as the relative mean intensity between them. We hypothesized that the nucleolus disassembly dynamics could be effectively treated as a quasi-equilibrium process during the slow phase of nucleolar disassembly. With this hypothesis, the two phases were in thermodynamic equilibrium at each time point of the experiment. We quantified the free energy of transfer (Δ*G*), which encapsulates the energy necessary for the transfer of a molecule from the dilute to the dense phase (32) (Fig. 2E, Supplemental Discussion). This analysis showed that the free energy remained negative and constant for cells in interphase, consistent with the observed stability of nucleolus during this stage. In contrast, as cells approached mitosis, Δ*G* gradually became less negative, turning positive around the time of the nuclear envelope breakdown. Collectively, these results demonstrate a continuous loss of thermodynamic stability in the nucleolus during the slow disassembly phase.

We next sought to examine the role of the slow phase in the overall disassembly of the nucleolus. The slow phase may prepare the nucleolus for the fast phase to occur at nuclear envelope breakdown, or it may be dispensable for the overall disassembly of the nucleolus. To test this, we treated interphase cells with 0.1% Triton-X 100 to rapidly permeabilize the nuclear membrane during imaging. We found that while this treatment was able to extensively disrupt the nuclear envelope and caused a rapid reduction in the fluorescence intensity of free NPM1 in the nucleoplasm, yet the nucleolus was stable over a much longer timescale (Fig. 2F, Movie S3).

This result suggested that the biochemical events during the slow phase of disassembly are necessary precursors for the rapid disassembly of nucleolus during the second rapid phase.

Next, we interrupted the progression of cells towards nuclear membrane breakdown at late prophase using the CDK1 inhibitor RO3306 (35). We observed that this treatment indeed reversed chromosome condensation consistent with results from previous studies that the disassembly of nucleolus is controlled by CDK1 activity in the nucleus (19, 20, 36) (Fig. 2G). Interestingly, we found that after this treatment during the slow phase of disassembly, the slope index α stalled and increased back to the levels of a pre-disassembled nucleolus (Fig. 2G, Movie S4). Together, these findings demonstrate that the biochemical events during the slow phase of the nucleolus disassembly are reversible and involve a necessary, gradual, and continuous modulation of binding kinetics between nucleolar material. Thus, this stage prepares the nucleolus for the rapid, irreversible disassembly that occurs upon nuclear envelop breakdown.

### Binding affinities of nucleolar material decreases during the disassembly process

The nucleolus exhibits liquid-like properties characterized by rapid diffusion and exchange of material with the nucleoplasm (26, 32). Therefore, changes in the binding kinetics during disassembly may alter the diffusivity of nucleolar material. To test this hypothesis, we performed fluorescence recovery after photobleaching (FRAP) experiments on individual nucleoli (Figs. 3A, S5A, S5B, Movie S5). For cells in interphase or early prophase, the fluorescence signal in a bleach spot with a diameter of 1 μm recovered to the pre-bleach level in less than one minute, indicating that the NPM1 protein was highly dynamic and rapidly exchanging with the pool of free NPM1 in the nucleoplasm, consistent with previous reports (19). We hypothesized that the biochemical changes of nucleolus during the slow phase of disassembly (Fig. 2C) might affect the NPM1 mobility. To explore this, we followed individual nucleolus along the disassembly trajectory and performed FRAP on the same nucleolus as cells transitioned from prophase to mitosis. We found that the fluorescence recovery rate continuously increased and significantly accelerated within 10 minutes prior to nuclear envelope breakdown (Figs. 3A-3C). Specifically, the fluorescence half-recovery times t_1/2_ decreased by more than threefold from early to late prophase (Fig. 3B), corresponding to an increase of the effective NPM1 diffusion coefficient from 0.14± 0.02 μm^2^/s to 0.36 ±0.14 μm^2^/s (Fig. 3C). By contrast, we did not observe any significant change in NPM1 diffusivity when we performed similar experiments on interphase cells (Figs. 3B, 3C). Interestingly, the effective diffusion coefficient for NPM1 is two orders of magnitude smaller than for unconjugated-GFP molecule in the nucleolus (25 μm^2^/s) (37), suggesting that NPM1 diffusion is significantly retarded by binding interactions with other nucleolar components.

**Figure 3.**
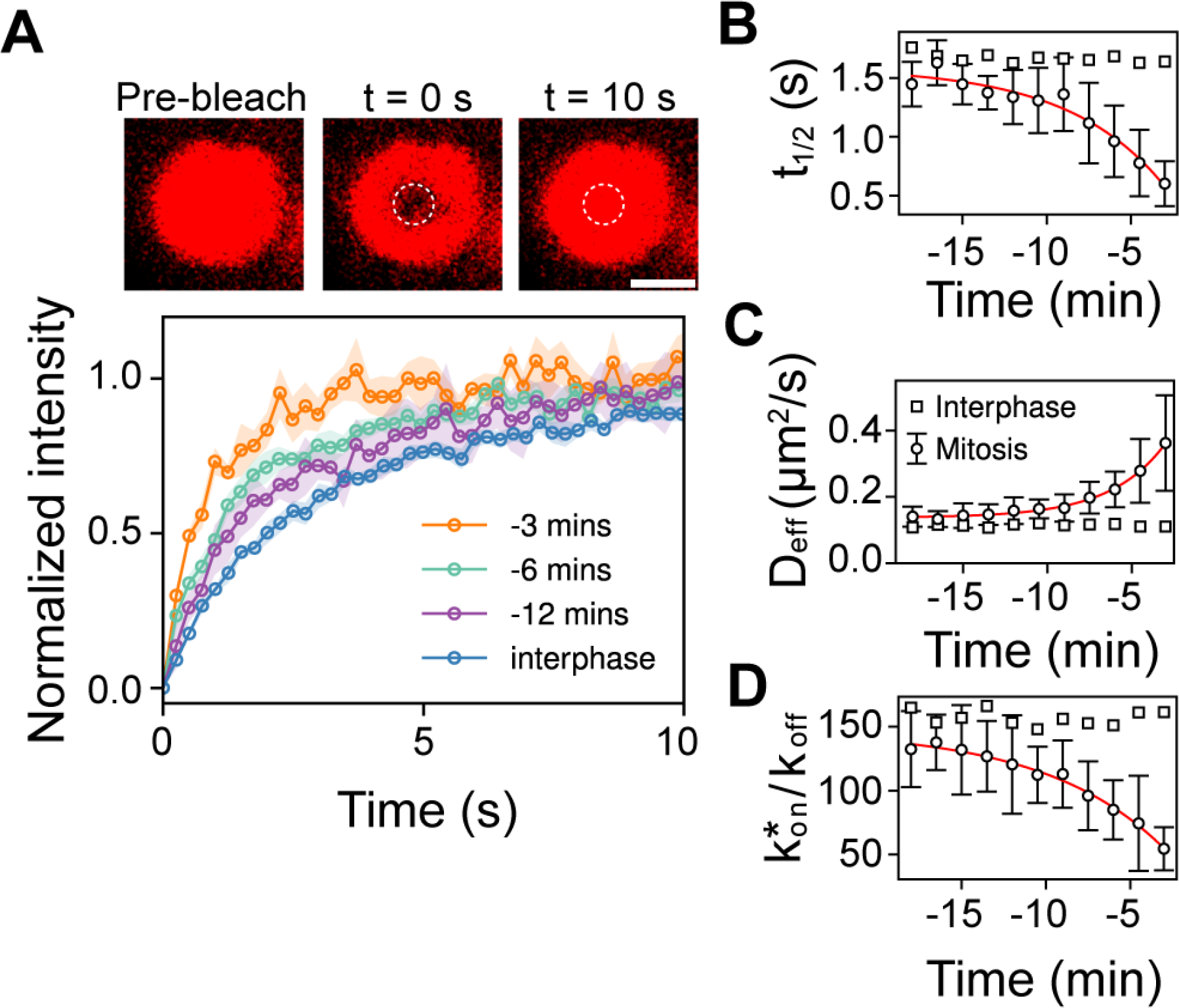
Nucleolus disassembly during prophase results in decreased of NPM1 binding kinetics. **(A)** Top, FRAP time course of nucleolus in HeLa cell expressing mRuby3-NPM1. Dashed circles indicate the bleached spot. Scale bar, 2μm. Bottom, quantification of the mean mRuby3-NPM1 fluorescence intensity recovery after FRAP at different time points of nucleoli through the prophase, and for cells in interphase (blue). (**B**) The fluorescence half-recovery times t_1/2_ quantified at different time points. (**C**) The effective diffusion coefficient of mRuby3-NPM1 at different time points. (**D**) Relative association/dissociation rate of NPM1 calculated from FRAP experiments. Red solid lines represent a single exponential decay function (B, D) and growth (C). Plot markers represent mean value, shaded areas (a, n=8 cells) and error bars (B-D, n=20 cells) represent mean ± s.d.

Next, we quantitatively estimated the binding kinetics by measuring the association and dissociation rates of NPM1, through fitting FRAP recovery curves with an effective diffusion-binding model reported previously (38) (Supplemental Discussion, Figs. S5A, S5B). We found that the relative binding constant of nucleolar material, 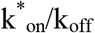, decreased along the disassembly trajectory of nucleolus during prophase while remaining constant for cells in interphase (Fig. 3D). Together, these results show that the process of nucleolus disassembly is a continuous transition rather than an abrupt and stepwise transition. We propose that this continuous modulation in the binding kinetics of nucleolar material during the disassembly of the nucleolus is a probable mechanism underlying the tuning of bulk material properties of the nucleolus and the morphological modifications that we have characterized (Figs. 1F, 1G). Furthermore, these changes prepare the nucleolus for complete disassembly at the nuclear envelope breakdown so that the disassembly process could be synchronized with the cell cycle.

### Material properties of nucleolus change during the disassembly process

The modulation of binding kinetics between nucleolar material throughout prophase (Fig. 3) that happens at the microscopic scale manifests the morphological changes of the nucleolus (Fig. 1), and potentially bulk material properties at the macroscopic scale. To further quantify this, we measured the bulk material properties of nucleolus during disassembly. Previous work to measure the surface tension of nucleolus relied on simplified assumptions (39). We adopted and optimized a methodology based on power spectrum analysis of nucleolus and nucleoplasm interface fluctuations (40, 41) (Figs. 4A, 4B, Fig. S6) (Supplemental Discussion, Movie S6). The significant advantage of this noninvasive analysis is that we were able to directly measure the surface tension (σ) and bending rigidity (κ) of the nucleolus from the shape fluctuation spectra alone. This analysis showed that the overall amplitude of the shape fluctuations, <|u(q)|^2^>, of the nucleolar interface increased as cells approached nuclear envelope breakdown (Fig. 4C). At each time point, the experimental amplitude spectra follow two power law regimes (*q*^γ^) for small and large q modes. The small q regime is dominated by surface tension contribution with an exponent of γ=-1 (*q*^-1^), and the large q regime is dominated by bending stiffness with an exponent γ=-3 (*q*^-3^). Strikingly, these measured exponents agree with the predictions of the theory vesicular membrane fluctuations (40-42) (Fig. 4C). We performed a global least-squares fit of our theoretical model to the experimental spectra (42) and determined the surface tension (μ) and bending rigidity (κ). In the bending rigidity dominant regime, the bending rigidity coefficient κ was obtained by fitting the data into theoretical model, while disregarding the surface tension (σ=0) (Supplemental Discussion, Fig. S7). Subsequently, in the surface tension dominant regime, the experimental data was fitted with κ as a constant, and σ as a single fit parameter (Supplemental Discussion, Fig. S7). We found that both μ and κ decreased nonlinearly as cells progressed towards the nuclear envelope breakdown (Fig. 4D-E). Interestingly, the macroscopic parameters characterizing the bulk material properties of nucleolus displayed a qualitatively similar variation pattern during disassembly to the one observed with the microscopic parameters (Fig. 3D). Thus, surface tension and bending rigidity serve as practical parameters in quantifying the collective impact of complex biochemical processes during nucleolus disassembly. Genetic manipulations, or pharmacological treatments could alter these parameters to disrupt nucleolar function (23).

**Figure 4.**
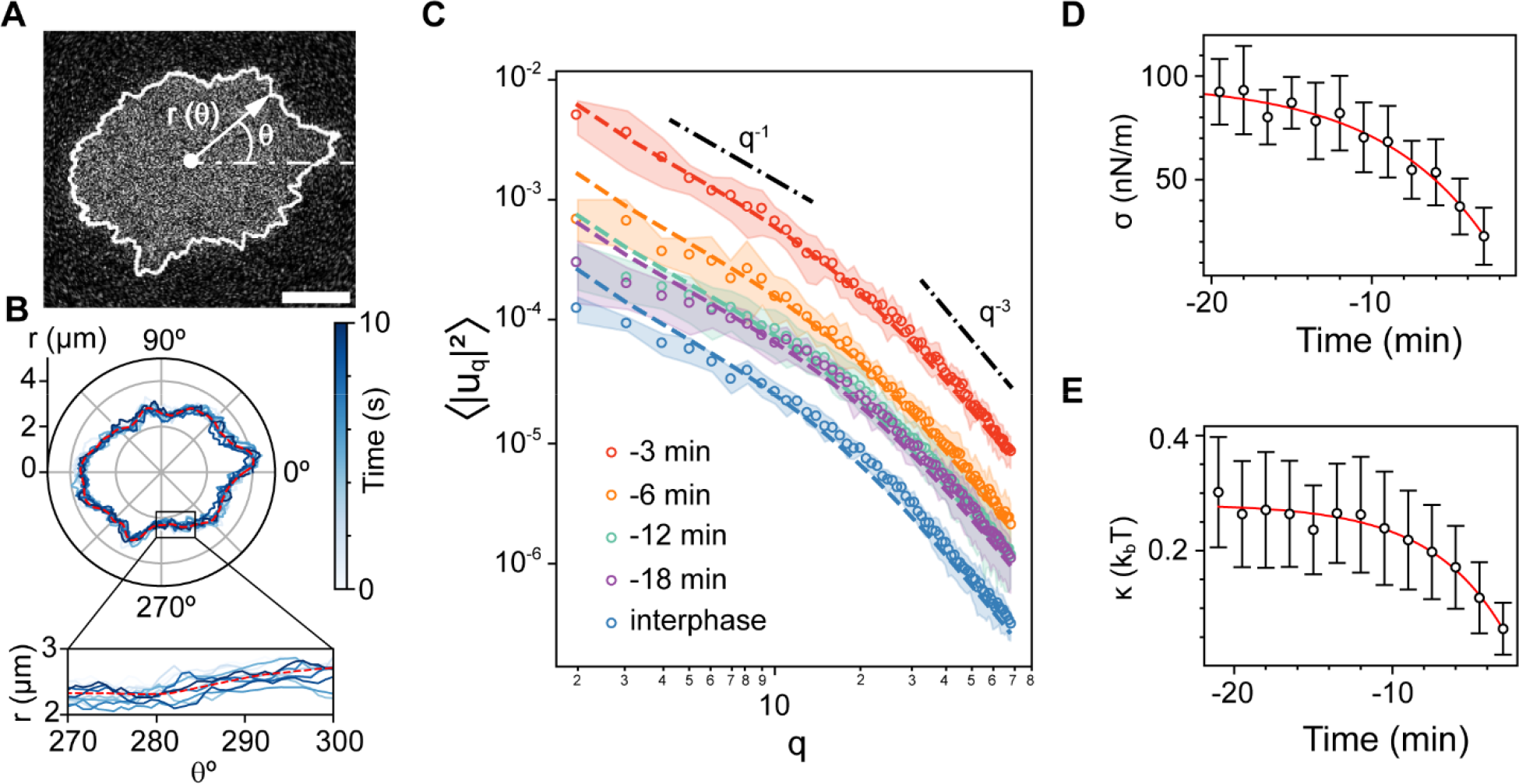
Reduction of nucleolar material properties during the disassembly process. **(A)** Representative image of a HeLa cell nucleolus. The white contour denotes the nucleolar shape. The white dot is nucleolar center of mass and the origin of polar coordinate. Scale bar, 2 μm. (**B**) Contours of nucleolus plotted in the polar coordinates at different time points for a cell in prophase denoting the shape fluctuation around a mean contour (red dashed line). Inset represents a zoom-in view. Color bar shows the time indicator for contours in the time series. (**C**) Shape fluctuation spectra as a function of mode number at different time points before nuclear membrane breakdown, and for cells in interphase (blue). Plot markers represent mean value, and shaded areas represent 95% bootstrap confidence intervals of standard deviation from n=10 cells. Dashed lines are the best fit data for experimental results from the theoretical model. Dash-dot lines represent surface tension dominant regime (q^-1^) and bending rigidity dominant regime (q^-3^), respectively. Surface tension (**D**) and bending rigidity (**E**) obtained from fitting experimental results with theoretical model as shown in (C). Red solid lines represent a single exponential decay function fit. Plot markers represent mean value, and error bars represent 95% bootstrap confidence intervals of standard deviation with n=22 cells.

### A critical regime in nucleolar disassembly

Previous results demonstrated a covariance between microscopic nucleolar features and macroscopic properties during the disassembly process (Fig. 3D and Fig. 4D). To connect these properties, we reduced the complex microscopic details into the effective physical parameter of NPM1 interaction strength with nucleolar material, defined as 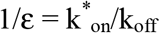. Notably, as cells progressed towards nuclear membrane breakdown the interaction strength of NPM1 with nucleolar components decreased resulting in an increase of ε (Fig. 5A). By comparing the variation of the surface tension (σ) (Fig. 4D) and the interaction strength (ε) (Fig. 3D) we found evidence of a power-law scaling in the standard form as σ=σ_o_(ε_c_ - ε)^μ^, where ε_c_ is the reciprocal of the interaction strength at critical point, μ_o_ is critical surface tension, and μ is critical exponent (Fig. 5B). We fitted the data to a series of power laws with μ = 0.92 to μ =1.35 to estimate ε_c_ (Fig. 5B, Supplemental Discussion). This interval was determined based on a qualitative assessment of the data and the scaling exponents for 2D (μ =1) and 3D (μ =1.26) Ising universality (28). We next determined how far the proposed critical regime extended in our system by plotting a normalized surface tension (σ/σ_o_) with respect to a normalized interaction strength difference (ε_c_ - ε)/ε_c_ (Fig. 5C). Based on this we found evidence that the power-law scaling extended up to 70% of the critical point ε_c_, which represents a significantly wider critical regime than previous predictions using numerical approaches (30, 43). Interestingly, the surface tension of the nucleolus during interphase does not fall within the critical scaling (Fig. 5C). Within the framework of critical phenomena, our data strongly suggests that cells exhibit an incredible capacity to tune microscopic interactions. These adjustments translate into changes in nucleolar surface tension at macroscopic scales of the entire condensate, ensuring robust and quantitative regulation of nucleolar disassembly during mitosis or in response to signaling cues.

**Figure 5.**
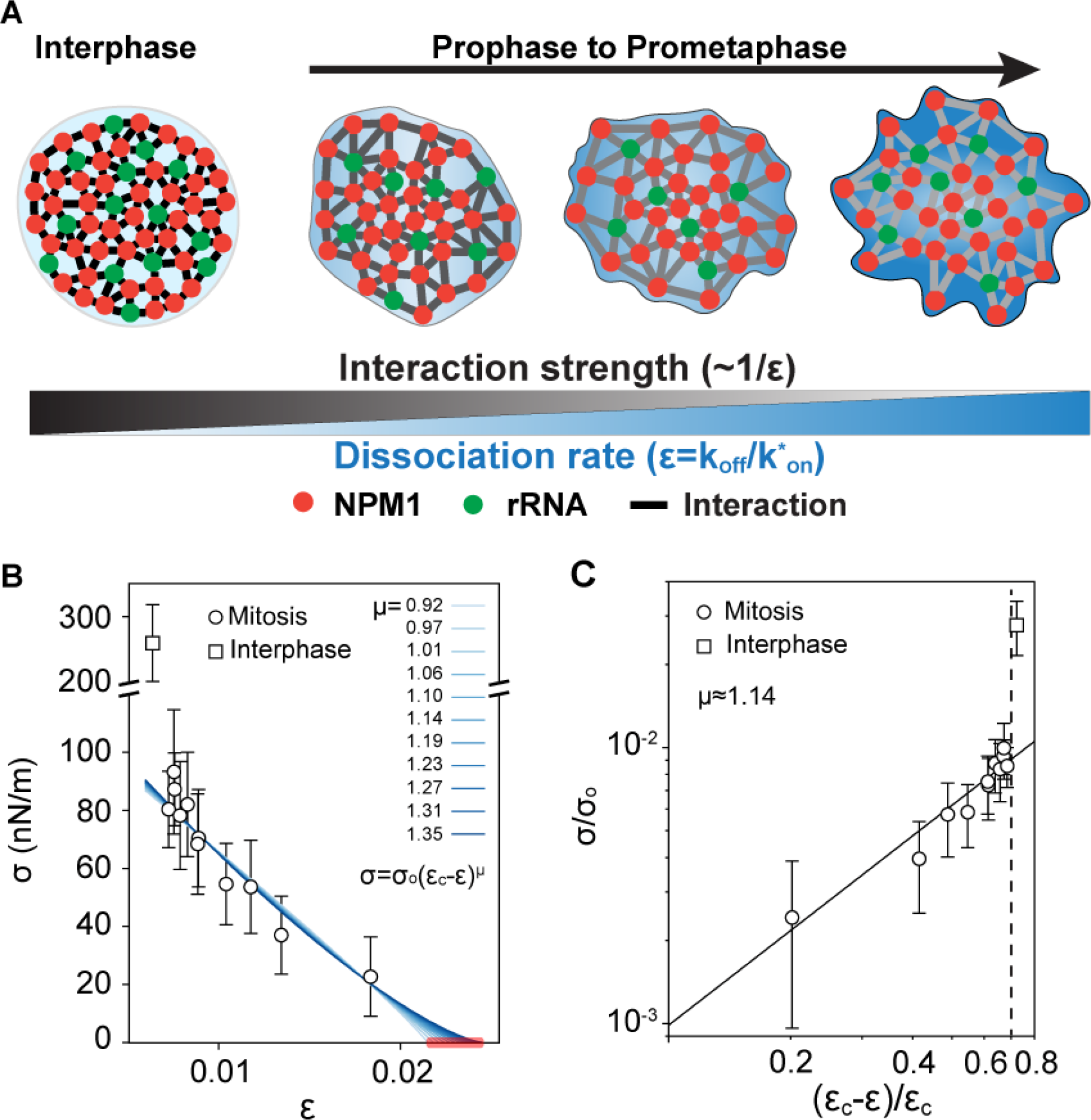
Range of the critical regime. (**A**) A schematic describing the emergence of nucleolar morphologies and surface tension changes at macroscopic scale as cells regulate the NPM1 interaction strength at microscopic scale throughout nucleolus disassembly. (**B**) Surface tension fitted to power law function with different critical exponent μ resulting in the critical parameter ε_c_ in a range from 0.0215 to 0.0245 (red bar). (**C**) Reduced surface tension (σ/σ_o_) as a function of the normalized interaction strength difference ((ε_c_-ε)/ε_c_) in a log-log scale. Solid line is power-law function with the critical exponent of μ=1.14. Vertical dashed line indicates estimated point where the system deviates from critical scaling law. error bars represent 95% bootstrap confidence intervals of standard deviation with n=22 cells.

## Discussion

Biomolecular condensates provide a general mechanism for cellular organization and compartmentalization. While the relation between the morphology and function of membrane-bound organelles has been established (7, 44, 45), the correlation between alterations in condensate morphology, material properties, and underlying microscopic changes remain unexplored. Our findings show that disassembly of the nucleolus in mammalian cells during mitosis is a continuous process that proceeds in two phases with distinct rates. The first phase starts in prophase and proceeds slowly for about 30 minutes. Interestingly, the changes that we characterized in this phase are reversible, and pharmacological inhibition of CDK1 is enough to restore interphase morphology. During the first phase, the free energy of transfer increases and binding kinetics of nucleolar material decreases, indicating that nucleoli are experiencing structural instability. Therefore, it is likely that the slow disassembly phase is driven by the continuous modulation of interaction strength between nucleolar materials. Within ten minutes before nuclear breakdown, the disassembly accelerates rapidly until the nucleolus disappears along with the nuclear envelope breakdown, likely due to rapid dilution. The two-step design, with a gradual and reversible preparatory phase preceding a rapid phase concurrent with nuclear envelope breakdown, enables the preservation of nucleolar structural stability while synchronizing its dynamics with the cell cycle. Our quantitative modeling demonstrated that the complex interplay of various biochemical events occurring within the nucleolus at the microscopic scale can be effectively probed at the macroscopic scale. Specifically, we showed that changes in the interaction strength of NPM1 with other nucleolar components during mitosis can modulate nucleolar surface tension changes at the macroscopic scale of the condensate. We provide evidence for a scale invariance within the dynamics of nucleolus disassembly near mitosis. Due to the limitations in manipulating molecular interactions under physiological conditions, we lack sufficient data to numerically establish a precise value for the critical exponent. However, we observed a wide parameter regime within which a power-law dependence between microscopic and macroscopic parameters of the condensate is maintained. Beyond this regime, nucleolar surface tension significantly deviated from the scaling behavior. This quantitative analysis provides compelling evidence for a nonequilibrium example of a critical phenomenon in biology. The analysis and conceptual framework developed here could be generally applied to study the structural dynamics of many other condensates in cells.

## Acknowledgements

This work was supported by the National Institutes of Health grants R01GM140272 (to R.V.), R01GM149076 (to X.W.) and the National Science Foundation-Simons Center for Quantitative Biology at Northwestern University and the Simons Foundation grant 597491 (to M.M.) and by The Searle Leadership Fund for the Life Sciences at Northwestern University (to R.V.). M.M. is a Simons Foundation Investigator.

## Materials and Methods

### Cell cultures, cell lines and transfections

Human HeLa S3 cells (ATCC CCL-2.2) were cultured in high glucose Dulbecco’s modified Eagle’s (DMEM) (Corning) medium supplemented with 10% fetal bovine serum (FBS), 1% (v/v) Penicillin/Streptomycin, 1% (v/v) Glutamax 100X, and 10 mM HEPES (pH 7.4, Gibco). Imaging media had similar formulation with cell cultured media without phenol red to reduce autofluorescence. Cells were grown in T25 tissue culture flask (Fisher Scientific) pre-coated with 0.1% (w/v) gelatin solution (Sigma) and maintained at 37 °C in a humidified atmosphere with 5% CO2. Cells were trypsinized (Trypsin, 0.25%) and passaged when they reached 80% confluency. Chromatin was visualized by stable expression of histone H2B fusion with mClover3. Lamin-AC was visualized by stable expression of fusion with mRuby2. NPM1 protein was visualized by tagging with mRuby3 or EGFP.

Hela cell line stably expressing mClover3-H2B and mRuby3-NPM1 were generated by PiggyBac transposons as previously described (*1*). In brief, all cDNA-expressing vectors for stable transfection were constructed as PiggyBac (PB) vectors with the CAG promoter to express genes of interest and PGK promoter for a drug resistance gene. PB-mClover3-H2B::PGK-BSD and PB-mRuby3-NPM1::PGK-Puro were co-transfected with CAG-PBase into HeLa S3 using a FUGENE HD transfection kit (Promega). Stable clones were obtained by sequential Puromycin and blasticidin selection. Clones with uniform expression of both mClover3-H2B and mRuby3-NPM1 were chosen for subsequent analysis.

HeLa cell line stably expressing EGFP-NPM1 and mRuby2-Lamin A/C were generated by lentiviruses transduction, encoding fluorescently tagged proteins of interest. Separate lentivectors containing EGFP-NPM1 (Addgene plasmid # 17578) and mRuby2-Lamin A/C (Addgene plasmid # 55901) were produced in HEK293T cells. Briefly, HEK293T cells were transfected with transfer plasmid and two helper plasmids psPAX2 (Addgene plasmid #12260) and VSV-G envelope expressing plasmid (pMD2.G, Addgene #12259) with Lipofectamine-3000 (Invitrogen). Supernatant was collected 2–3 days post-transfection, cell debris was pelleted by centrifugation (3000xg for 5 minutes) and discarded and the supernatant containing virus was kept at 4 degree until use. One day before transduction, target HeLa cells were seeded in 24-well plate at 40% confluency per well. The next day, the cell was transduced with lentivirus-containing EGFP-NPM1. 48 h after transduction, cells were trypsinized and moved into a T25 flask to grow before sorting. The midrange expressing cells were kept after sorting. In the second stage, the stably expressing EGFP-NPM1 Hela cells were transduced with lentivirus-containing mRuby2-Lamin A/C using the same procedure. Next, cells were dual sorted for collecting populations that expressed EGFP-NPM1 and mRuby2-Lamin A/C. Sorted cells were grown in a large T125 flask and were aliquoted and stored in liquid nitrogen for future use.

### Microscopy and sample preparation

Cells were seeded into a 4 chamber-35 mm glass bottom dish (Cellvis, *D35C4-20-1*.*5-N*) pre-coated with 1% type I bovine atelocollagen (PureCol, 5005) at 35% confluency and imaged after 24h at 70-80% confluency. 30 minutes before imaging, the media was changed to the pre-warm imaging media. During live cell imaging the cells were in the environmental chamber at 37ºC, high humidity with 5% CO_2_. All images were recorded on a Zeiss LSM 800 laser scanning confocal microscope, using a 40x/1.3 NA oil immersion objective, operated by ZEN Blue 2011 software. A 488nm laser was used to image mClover3-H2B and EGFP-NPM1 proteins. A 561nm laser for imaging mRuby3-NPM1 and mRuby2-Lamin A/C proteins. For time-lapse imaging, a volumetric image of 551pixel x 551pixel x 30 z-stack (40 _μ_m x 40 _μ_m x 30 _μ_m) was taken every minute with an exposure time of 160 ms. This time-lapse image ended 5-10 minutes after nuclear envelop breakdown.

### Cell treatments with RO-3306 and Triton X-100

RO-3306 was dissolved in DMSO at 10 mM stock concentration (Sigma). For RO-3306 treatment, 2X working concentration of RO-3306 (20 _μ_M) was prepared in warm imaging media. First prophase cells were imaged for 20 minutes at framerate of 1 frame per minute. The imaging was then paused, and half of the media in the chamber was exchanged with the media containing RO-3306 to achieve a final working concentration of 10 _μ_M (*2*). The cells were recorded for an additional 1 hour immediately after the addition of RO-3306. This procedure was also used for interphase cells to treat with 0.1% Triton X-100 to permeabilize nuclear membrane.

### Fluorescence Recovery After Photobleaching (FRAP) imaging setup

FRAP experiments were done on the same microscope setup with excitation laser wavelength of 561 nm operating at 5% maximum laser power (10 mW). This power was 500-fold higher than the laser intensity used for image acquisition, which was high enough so that the bleach spot intensity was reduced up to 60%. 10 prebleach images (256pixel x 256pixel = 8 _μ_m x 8 _μ_m) were acquired, followed by 10 scan iterations of rapid photo-bleach (lasted ∼170 ms) in a spot of 1_μ_m in diameter. The bleach spots were chosen at the center of nucleolus to avoid any boundary effects. The post-bleach images were recorded at speed of 4 frames per second for one minute to ensure full recovery. FRAP experiment was repeated every 1.5 minute on the same nucleolus until nuclear membrane breakdown. See below for more details about FRAP data analysis.

### Imaging acquisition for measuring nucleolar surface fluctuations

We collected surface fluctuations of nucleolus using the same microscope setup described above with optimized image acquisition settings. The image field of view was fixed at 256pixel x 256pixel (8 _μ_m x 8 _μ_m). We acquired images continuously at equatorial plane of nucleolus with an exposure time of 57 ms, corresponding to a total of 173 images in 10 s (17.3 Hz). The acquisition repeated every 1.5 minute until nucleolar breakdown. The details of shape fluctuation analysis is described in supplementary text.

## Supplementary Figures

**Fig. S1.**
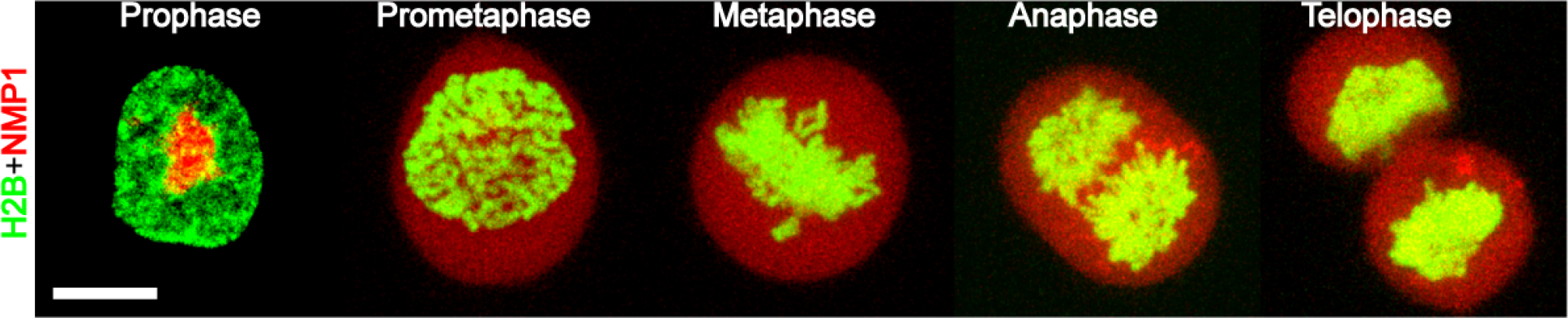
Fluorescent images of a HeLa cell expressing mClover3-H2B (green) and mRuby3-NPM1 (red) demonstrate normal phases of mitosis in our experiments. All prophase cells that were selected to follow the disassembly dynamic of nucleolus progressed through mitosis normally unless they were exposed to pharmacological treatments. Scale bar, 10 μm.

**Fig. S2.**
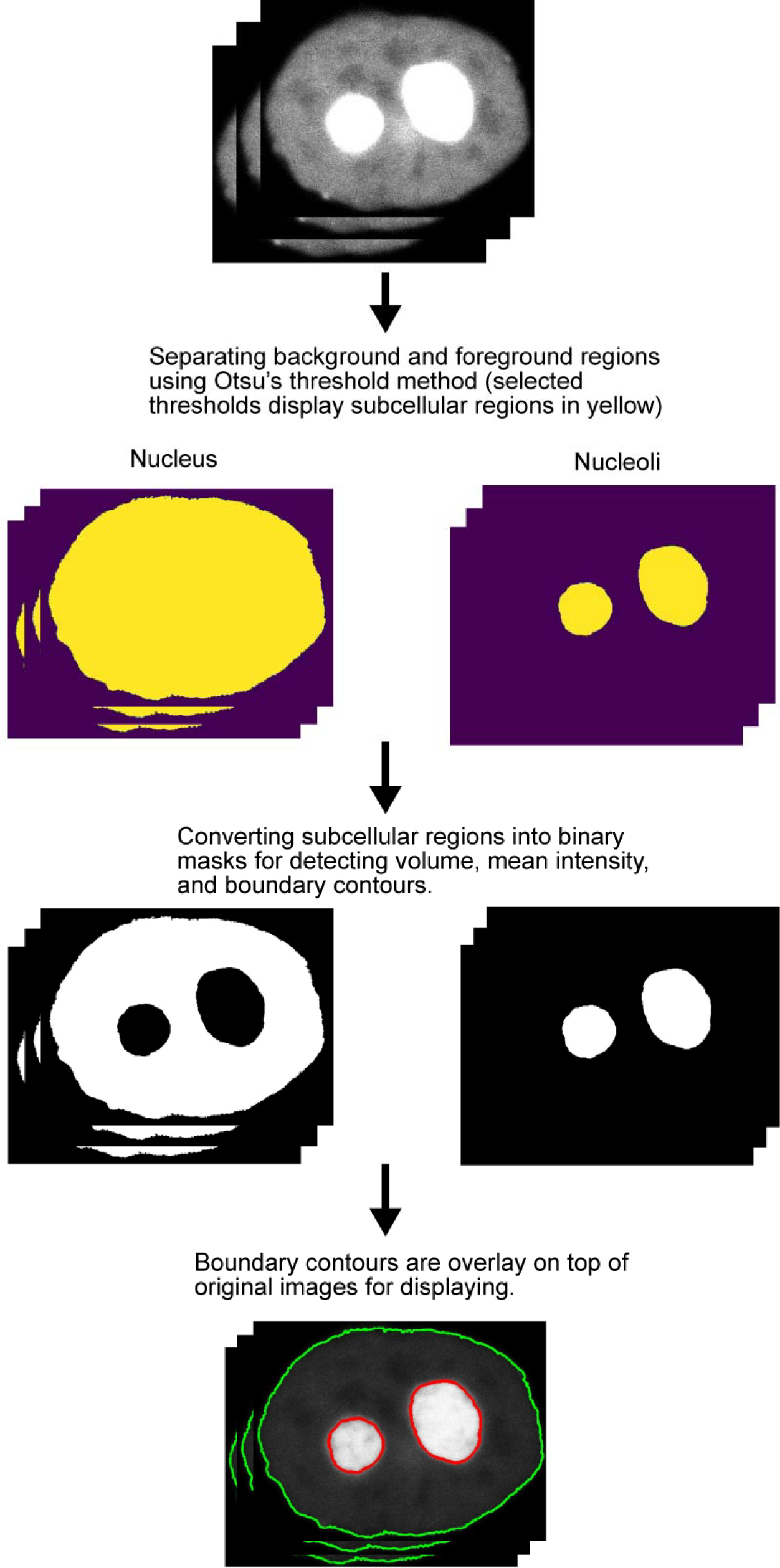
General steps for segmenting cell nuclear and nucleoli using Otsu’s thresholding method.

**Fig. S3.**
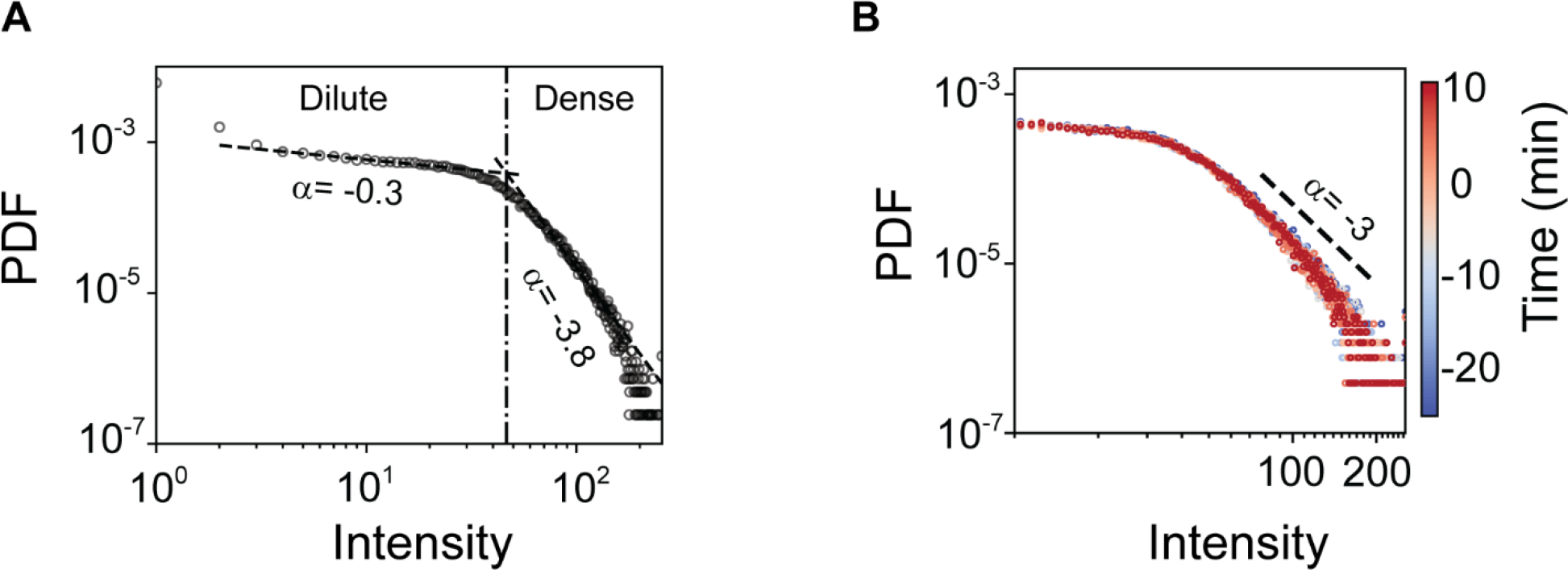
A. Probability density function of pixel intensity at single time point of a nucleolus in a cell at mitosis. Dashed lines represent the best fits into data using power law functions for dilute and dense phase, corresponding to the power law exponential of α=-0.3 and α=-3.8, respectively. **B**. Probability density function of pixel intensities at different time points at 1 minute time intervals of nucleolus at interphase, taking in the same experimental window with a cell at mitosis. t = 0 denotes the time of nuclear envelope breakdown. Dashed lines represent the power law exponent coefficients α of dense phase in log-log scale.

**Fig. S4.**
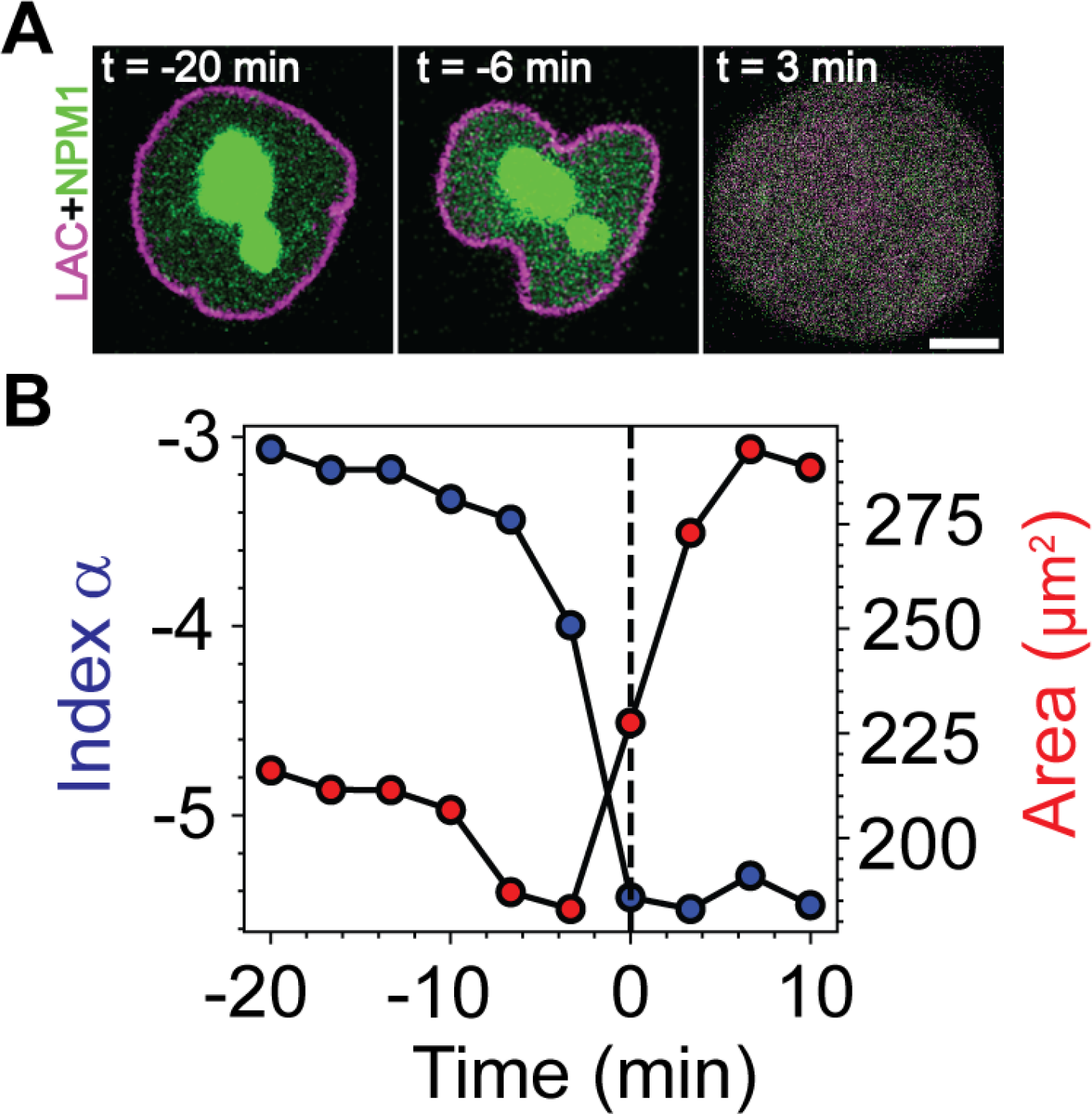
A. Representative images series of a HeLa cell expressing Lamin A/C-mRuby2 (magenta, to visualize nuclear membrane), and EGFP-NPM1 (green, to visualize nucleolus) progressing toward nuclear envelop breakdown. Scale bar, 5μm. **B**. Changes in the index _α_ (blue circles) and area enclosed by nuclear envelop (red circles) as a cell in prophase progressed toward nuclear envelop breakdown. Dashed line indicates t = 0 when nucleolus disappears and nuclear envelop breaks down.

**Fig. S5.**
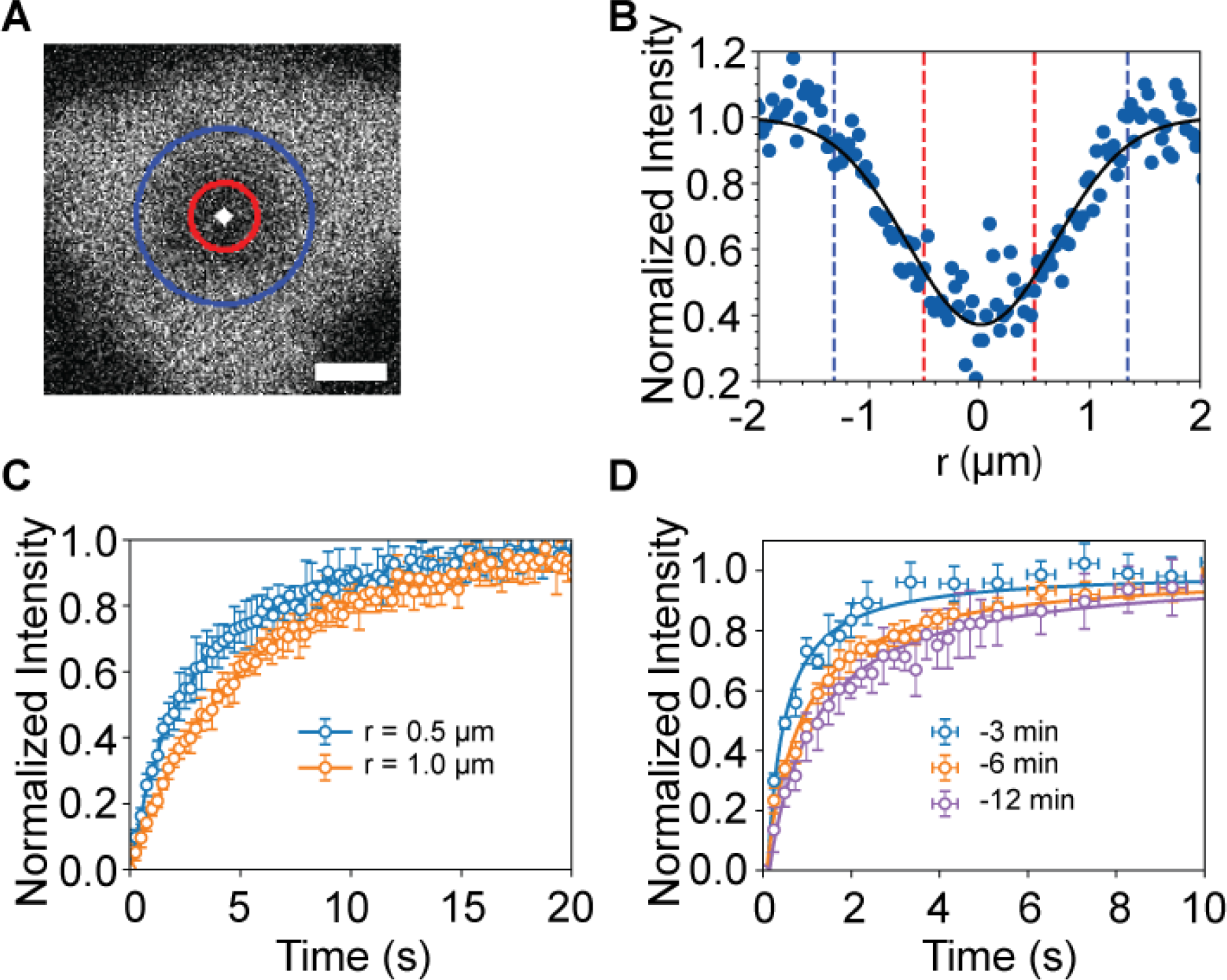
A. the initial post-bleach image of nucleolus after 10 times bleaching scans with 5% laser power. White dot, red circle, and blue circle represent center of the bleach spot, nominal bleach spot size (radius of r_n_=0.5 μm), and the experimental bleach spot with an effective radius of r_e_=1.35 μm, respectively. Scale bar, 1 μm. **B**. Fitting the initial post-bleach spot profile in (a) with Gaussian function (black solid line). **C**. FRAP curves show dependence of recovery rates on the size of the bleach spot. Plot markers represent mean values, and error bars represent mean ± s.d. with n=6 cells. **D**. FRAP data can be fitted into a single diffusion equation (solid lines). Plot markers represent mean values, and error bars represent mean ± s.d. with n=8 cells.

**Fig. S6.**
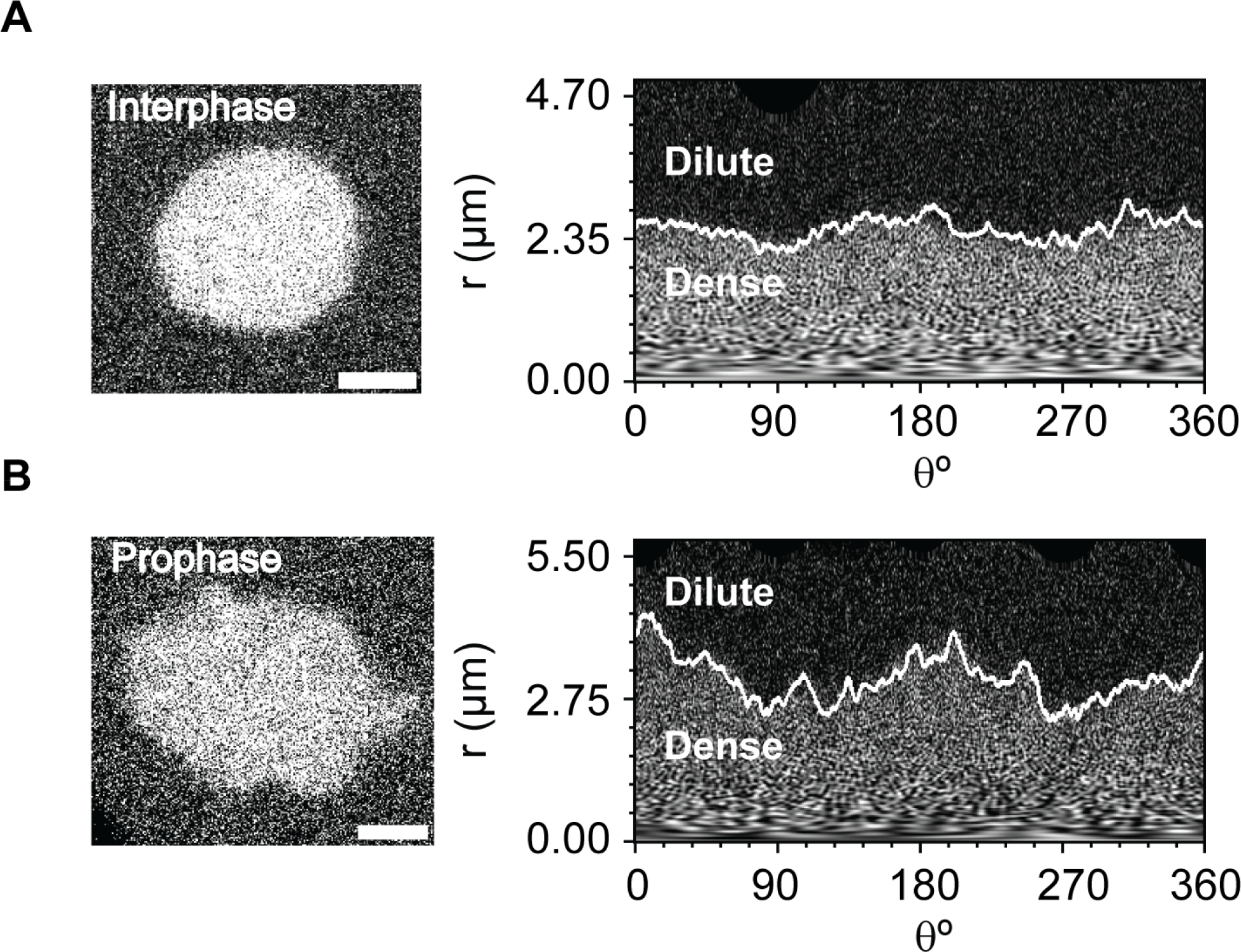
A-B. Left panels: a snapshot of nucleolus at interphase (A), and prophase (B). Right panels: interfacial boundaries between dense and dilute phases. White solid contours demonstrate the boundary. Horizontal and vertical axis represent angular (θ) and radial length (r) in polar coordinates, respectively. Scale bar, 2 μm.

**Fig. S7.**
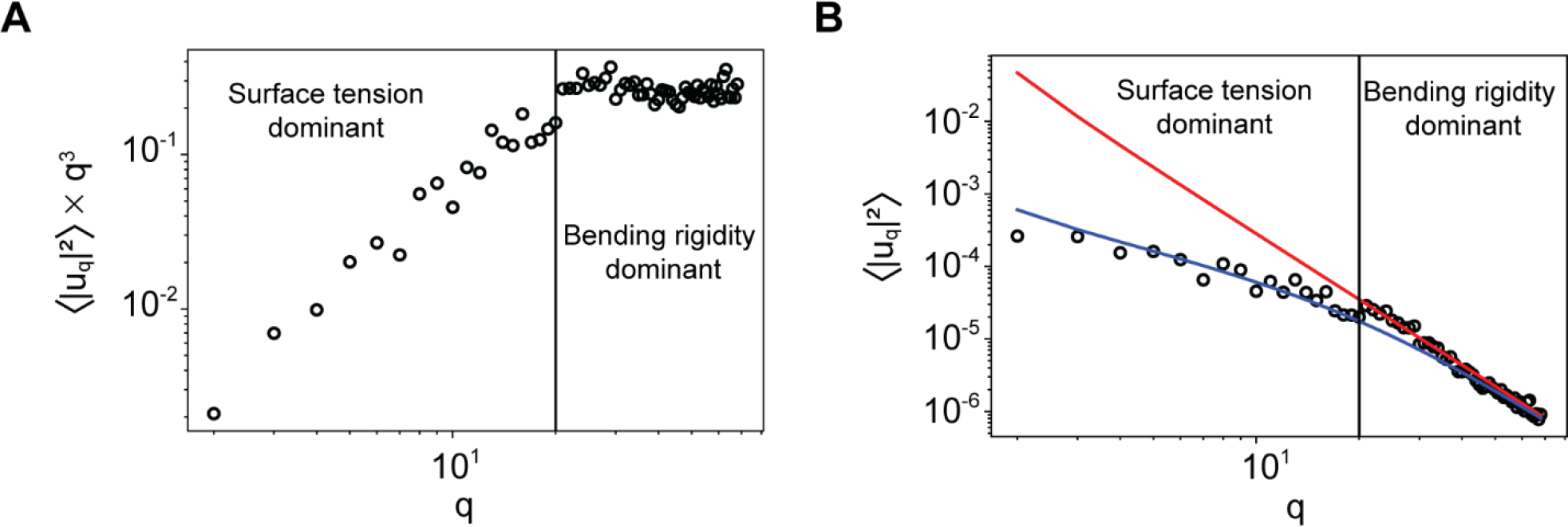
A. The plot of ⟨ lu_q_l^2^ ⟩ x q^3^ as a function the mode number q obtained from a nucleolus in a cell at mitosis. A plateau is observed at q values above a critical value (q_c_, black solid line) indicating bending rigidity dominant regime. **B**. Fitting results overlay on top of experimental data shown in (A), red solid line represents the fitting into bending rigidity dominant regime alone, while blue solid line is fitting result including the surface tension.

